# Comparison of Age-Related Decline and Behavioral Validity in C57BL/6 and CB6F1 Mice

**DOI:** 10.1101/2024.06.14.599036

**Authors:** Gerald Yu Liao, Christina Pettan-Brewer, Warren Ladiges

## Abstract

Variability in physical resilience to aging prompts a comprehensive examination of underlying mechanisms across organs and individuals. We conducted a detailed exploration of behavioral and physiological differences between C57BL/6 and CB6F1 mice across various age groups. In behavioral assays, B6 mice displayed superior performance in rotarod tasks but higher anxiety while CB6F1 mice exhibited a decline in short-term memory with age. Grip strength, long-term memory, and voluntary wheel running declined similarly with age in both strains. Examining physiological phenotypes, B6 mice exhibited lower body fat percentages across ages compared to CB6F1 mice, though cataract severity worsened with age in both strains. Analysis of cardiac functions revealed differences between strains, with worsening left ventricular hypertrophy and structural heart abnormalities with age in CB6F1 mice along with higher blood pressure than B6. Lesion scores showed an age-related increase in heart, kidney, and liver lesions in both strains, while lung lesions worsened with age only in CB6F1 mice. This study underscores the validity of behavioral assays and geropathology assessment in reflecting age-related decline and emphasizes the importance of considering strain specificity when using mouse models to study human aging.

## Introduction

Aging is a universal phenomenon characterized by a gradual decline in physiological function and an increased susceptibility to disease, posing significant challenges to human health. A complete understanding of the underlying mechanisms of aging is therefore paramount for the development of effective treatments against age-related diseases, which are major contributors to morbidity and mortality in humans (1). Given that between-individual and between-organ heterogeneity in health suggests variability in response to physical stress (2), the documentation of resilience across the lifespan, starting at a relatively young age in response to physical stress, offers valuable insights into individual health trajectories and the promotion of healthy aging.

However, the complexities inherent in human-based research, including ethical, social, and cost considerations, coupled with the entered human lifespan, necessitate alternative approaches to studying aging. Animal models have emerged as pragmatic tools for investigating age-related disease progression and treatment responses. While no single animal model can fully replicate all aspects of human aging, a thorough understanding of the characteristics of specific models and judicious interpretation of results can facilitate targeted investigations into critical facets of age-related diseases and their treatments (3).

Mice, due to their relatively short lifespan and modifiable genetic makeup, are among the most widely used animal models in aging research. Characterized by its widespread availability and well-documented genetic, behavioral, and cognitive background, the inbred C57BL/6 strain stands out as a prominent model in the study of aging and age-related diseases (1). In addition, a cross between C57BL/6 and Balb/c, designated as the CB6F1 mice and characterized by its more heterogeneous genetic background compared to the parenteral inbred strains, has also been validated (4l). Both are readily available from the National Institute on Aging Aged Rodent Colony. Nevertheless, few studies have assessed difference in age-related behavioral and cognitive behaviors between the two strains (5-6), and no studies have conducted a comprehensive examination of differences based on age-related behavioral, physiological, and geropathological differences.

In the present study, we address this gap by conducting a series of behavioral assays (i.e., rotarod performance, radial water tread maze, grip strength, open-field test, voluntary wheel running), physiological phenotypes (i.e., cataract severity, metabolic assessment, cardiac functions (left ventricular hypertrophy, cardiac reserve, ejection fraction, structural heart abnormalities), blood pressure), and geropathological assessments (i.e., heart, kidney, lung, liver) in C57BL/6 and CB6F1 male mice at different ages. By systematically analyzing various parameters, we aimed to elucidate the differences in aging trajectories between these strains and provide comparative insights as models of aging and age-related diseases.

## Materials and Methods

### Animals

CB6F1 (C57BL/6 X Balb/c F1 cross) and C57BL/6 male mice in age groups of 4, 12, 20, 28 months were obtained from the National Institute on Aging Aged Rodent Colony, contracted by Charles River, Inc. Mice were housed in a specific pathogen free mouse facility at the University of Washington (UW) main campus in Seattle, WA. The status of the room was monitored under the guidance of the Rodent Health Monitoring Program within the purview of the UW Department of Comparative Medicine. Mice were group housed, up to five per cage, and given nestlets (Ancare Corp, Bellmore, NY) for environmental enrichment. Mice were acclimated for two weeks before starting test procedures. All procedures were approved by the University of Washington IACUC (Animal Care and Use Committee).

### Behavioral Assays

#### Rotarod Performance

Agility, a measure of balance and coordination, was assessed using a Rotamax 4/8 rotating bar machine (Columbus Instruments, Inc) (5,7). The machine tested the ability of mice to maintain walking speed on a rotating bar, with individual lanes separated to prevent visual or physical interactions. Seven photobeams were embedded in each lane of the enclosure with a software recording the photobeam breaks during the task. Once the animal falls from the rotarod, the final team is recorded because of the absence in beam breaks. The initial speed was set to 0 RPM and gradually increased by 0.1 RPM/s over a 5-minute duration until 40 rpm, when all mice fell off and were detected by a sensor. The time in seconds was recorded for each mouse over three trials and the mean time of the three trials per mouse was recorded.

#### Memory Assessment

A radial water tread maze was used to assess short-term and long-term memory (8). The maze consisted of a circular basin with nine holes, eight decoys leading to dead ends, and one escape hole leading to a dark safety box equipped with a heating pad to simulate a standard mouse cage. The basin contained approximately one inch of water and an overhead light placed above the cage as an escape incentive. Mice spent a maximum of 3 minutes per trial in the maze, with those that were stationary for greater than 10 seconds brought back to the center of the apparatus. Mice had three trials in the maze for four consecutive days of training, followed by testing on the fifth day and retesting on the twelfth day.

#### Grip Strength Assessment

Forelimb strength, a measure of frailty in elderly humans was measured using a Grip Strength Meter (9-10). Each mouse was positioned horizontally with forepaws on a metal grip bar (Columbus Instruments, Inc), and the mouse was pulled back at a uniform rate until releasing the bar. The machine recorded the maximum force exerted by the mouse for a total of five trials. Mice were weighed on the test day and peak force was expressed relative to body weight to normalize grip strength measurements.

#### Voluntary Wheel Running Assessment

Total distance ran over three days was measured with a running wheel added to a standard mouse cage (11). Mice were individually housed in standard cages with a slanted running wheel wirelessly connected to a computer (Med Associates, Inc). There was a two-day acclimation period with the wheels locked and on the third day the wheel was unlocked, and data collection began. Running distances were continuously monitored over a 72-hour period with total distances ran every minute recorded in kilometers.

#### Anxiety Assessment

An open-field photobeam testing system (OFT) (Columbus Instruments, Inc) was employed to evaluate anxiety-related behaviors of mice in a novel environment as previously described (12). The apparatus simulates a standard mouse cage, featuring a clear rectangular container and infrared beams arranged in a grid pattern, three horizontally and four vertically. The open-field photobeam system was configured with two sets of infrared beams to measure both lateral and vertical activity. Beam breaks, which occurred when mice crossed an infrared beam, were counted for each activity. The data collected were subsequently categorized into two distinct zones: the central and peripheral areas of the container. This categorization allowed for the assessment of anxiety levels based on preference for exploring specific regions. Increased time spent in the central area suggested reduced anxiety, whereas a preference for peripheral regions suggested heightened anxiety. Each mouse was placed inside the testing container for a period of five minutes. This standard duration ensured consistent evaluation of anxiety-related behaviors and minimized potential habituation effects or stress-related responses.

### Physiological Phenotypes

#### Cataract Assessment

Recognized as a marker of visual function in previous studies (13-15), cataract formation was assessed using slit-lamp ophthalmoscopy. Both eyes of each mouse were examined and averaged to ensure comprehensive assessment of cataract progression. Based on lens opacity, the degree of cataract progression was graded on a scale from 0 to 4, with increments of 0.5. A score of 0 represented complete clarity of the lens, while a score of 4 indicated the presence of a mature cataract occupying the entire lens.

#### Metabolic Assessment

Changes in body composition, even in the absence of body weight changes, is a well-documented factor of aging (16). Body composition, including total body fat and lean mass, were assessed using noninvasive quantitative magnetic resonance (QMR) imaging (Echo Medical System).

#### Cardiac Function Assessment

Echocardiography, a non-invasive procedure, was employed to assess systolic and diastolic function in mice (17). The Siemans Acuson CV-70 system was utilized, employing standard imaging planes including M-mode, conventional, and Tissue Doppler imaging. Parameters such as left ventricular mass index (LVMI), left atrial (LA) dimension, end-diastolic and end-systolic dimensions, LV fractional shortening, Ea/Aa (diastolic function) measured by tissue Doppler imaging of the mitral annulus, and Myocardial Performance Index (MPI) were measured, as previous described (18). Additionally, blood pressure measurements were obtained and cross-correlated with echocardiographic functional parameters to provide comprehensive cardiac function assessment.

### Geropathology

Mice in all groups were decapitated after euthanasia via 5 min carbon dioxide inhalation approved by Institute/Center (IC) Animal Care and Use Committee (ACUC). Necropsy was performed after euthanasia with a postmortem time of 5 min. Sections of heart, lung, liver, and kidney were flash frozen and stored at -80 degrees Celsius. Sections of the same organs were also fixed in 10% buffered formalin for 48 h then blocked in paraffin wax through University of Washington Department of Comparative Medicine’s Histology Lab. Sections of 4 um fixed tissues were stained with hematoxylin and eosin for geropathology grading by two board certified veterinary pathologists in a blinded manner according to published guidelines (19). The geropathology scores were tabulated into a composite lesion score (CLS) for each of four major organs: heart, lungs, liver, and kidney, and recorded for each animal. The CLS was calculated by adding the severity score (from 1 to 4) of each lesion in an organ for each mouse in a cohort, and then adding the total lesion score of each organ for all mice in a specific cohort and dividing by the number of mice in that cohort. Therefore, the CLS is a standardized score that can be used statistically to compare lesion severity among different cohorts.

### Statistical Analysis

All data was grouped according to strain and age. A Shapiro-Wilk test was used to assess data under each group to determine whether there was a normal distribution. Two-tailed student’s t-test was used to compare between results from each treatment cohort for normally distributed data while the Mann-Whitney U test was used to determine significant differences between two groups when the data was not normally distributed. For geropathology, scoring data was the average lesion score from two pathologists analyzed by cohort using the two-tailed student’s t-test. Mean values with standard error bars (SEM) are presented by group in the figures. Quantitative interval outcomes collected from both pathologists were normalized for central tendency analysis. Correlation analysis between treatment cohorts, strains and lesion scores was done by one and two-way ANOVA. All statistical tests were conducted at 0.05 significance level. All statistical analyses were performed using GraphPad prism (version 10.0.3).

## Results

### Rotarod performance was age-dependent, with B6 mice performing better

Older mice (20 to 28 months) performed worse compared to younger mice (4 to 12 months) in both strains (p’s < 0.01) (Figures 1A & 1B). Additionally, B6 mice consistently outperformed CB6F1 mice (Figure 1C) (p’s < 0.01).

**Figure 1.**
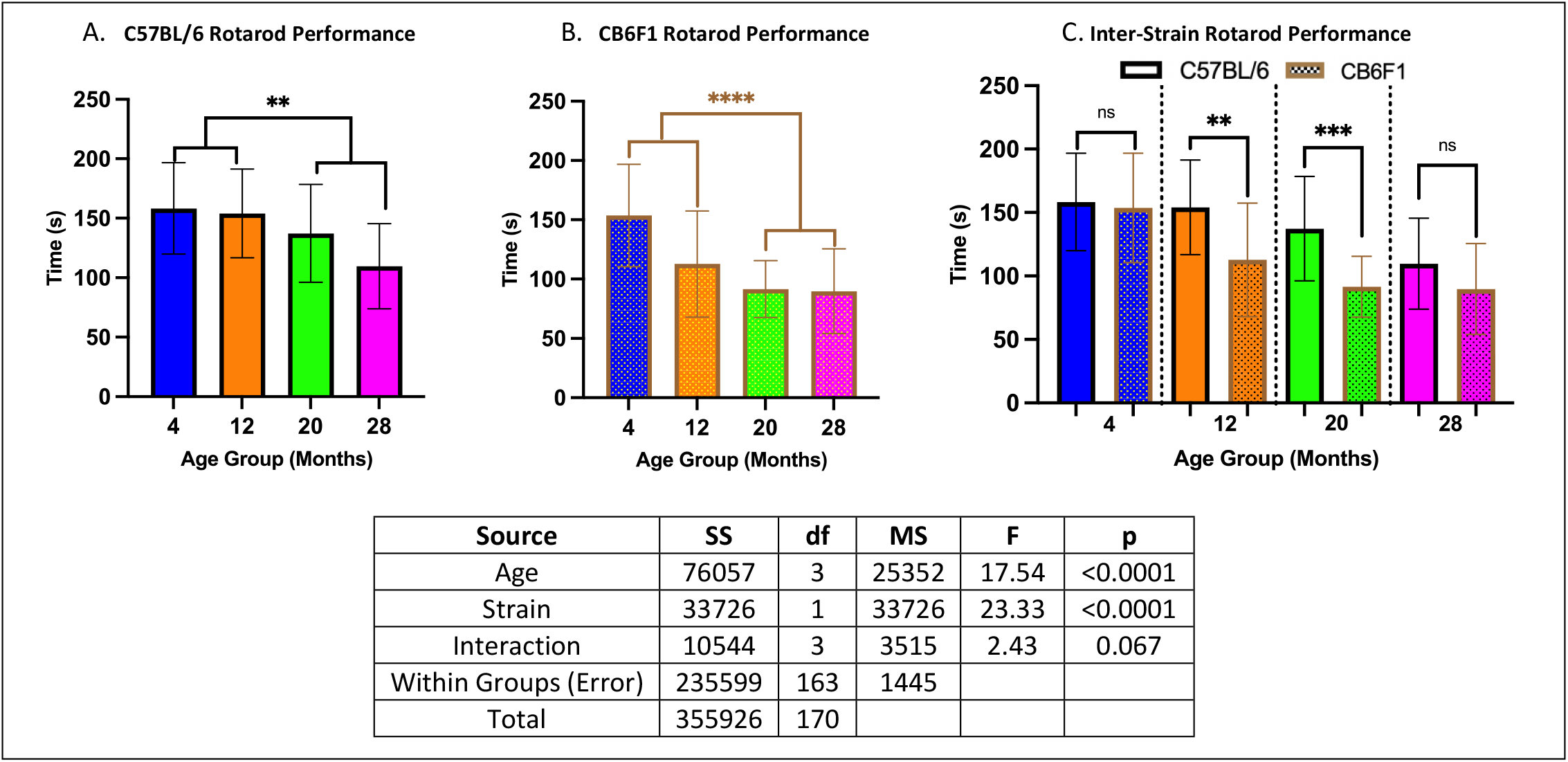
Rotarod Performance. **A**. CB6F1 mice at 20 and 28 months of age exhibited significant differences when compared to 4 and 12-month-old cohorts and showed a negative association between performance and age. **B**. B6 mice at the same age groups also showed statistically significant age-related decrease in performance as seen in CB6F1 mice. **C**. Inter-strain comparisons across age groups demonstrated greater performance in B6 compared to CB6F1 mice (** p < 0.01, ***p < 0.001, ****p < 0.0001, ns = not significant (p > 0.05), N = 19-29/cohort).

### Memory in the radial water tread maze was age- and strain-dependent

In C57BL/6 (B6) mice, memory performance did not differ significantly between older mice (20 to 28 months) and younger mice (4 to 12 months) (p’s > 0.05) (<u>STM</u>: Figure 2A, <u>LTM</u>: Figure 2D). However, in CB6F1 mice, older cohorts exhibited impaired memory compared to younger cohorts (p’s < 0.05) (<u>STM</u>: Figure 2B, <u>LTM</u>: Figure 2E). Additionally, B6 mice generally demonstrated better STM compared to CB6F1 mice (p’s < 0.05) (Figure 2C), while no significant differences were observed in LTM between the strains (p’s > 0.05) (Figure 2F).

**Figure 2.**
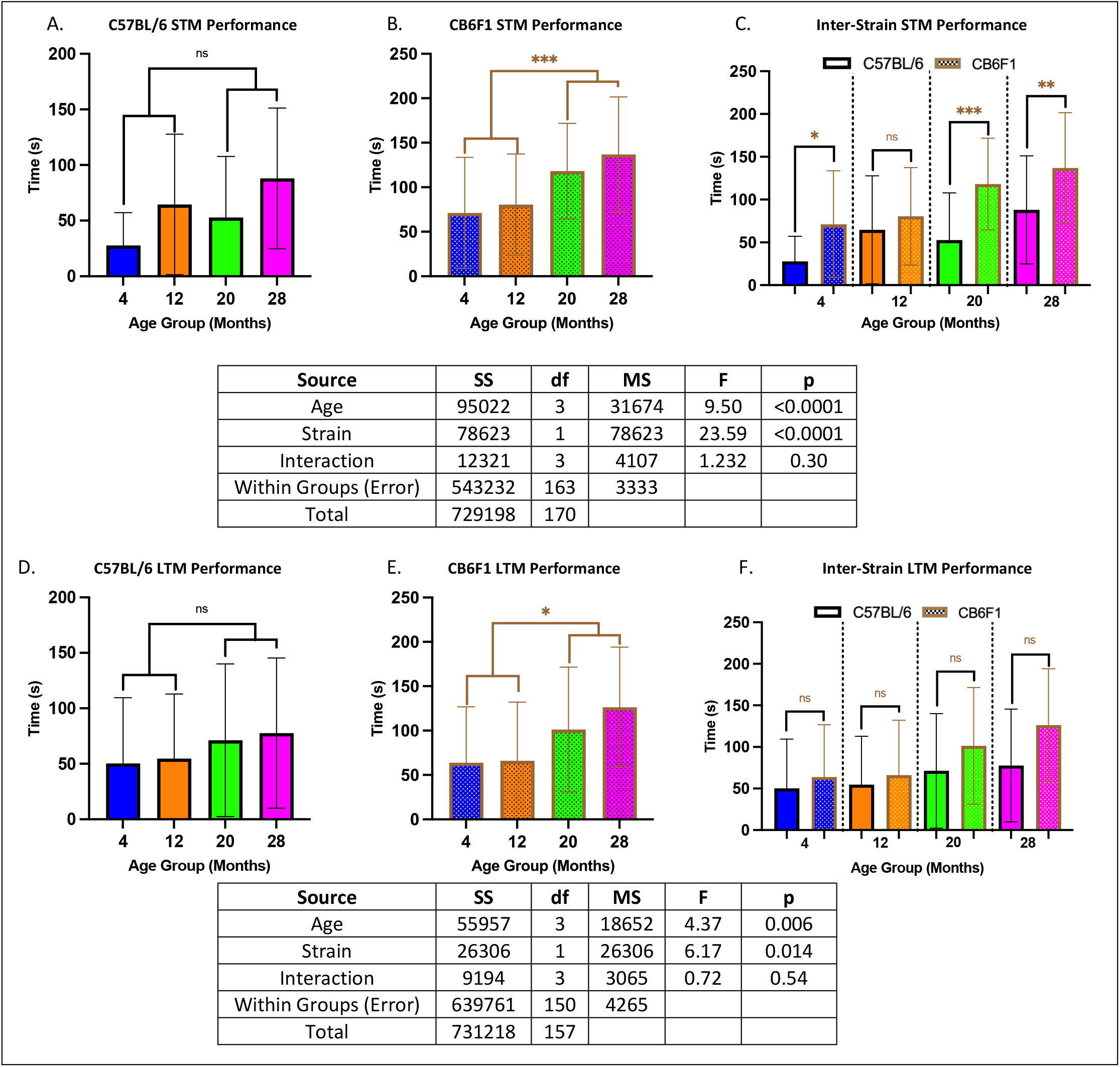
Memory Performance. **A/B. STM:** B6 mice at 20 and 28 months of age did not exhibit significant differences when compared to their younger counterparts, while CB6F1 mice at 20 and 28 months of age exhibited significant differences when compared to their younger counterparts. **C. STM:** B6 mice performed better than CB6F1 across age groups. **D/E. LTM:** Similar to STM, B6 mice did not exhibit age-related differences while CB6F1 did. **F. LTM:** B6 mice performed similarly to CB6F1 across age groups (*p < 0.05, **p < 0.01, ***p < 0.001, ns = not significant (p > 0.05), N = 19-29/cohort).

### Grip strength & wheel running performance decreased with age in both CB6F1 & B6 mice

Across all age groups, both strains exhibited similar performance, with no significant differences detected between them (p’s > 0.05). Specifically, grip strength was significantly lower in older cohorts compared to younger ones (p’s < 0.05) (Figures 3A & 3B). Similarly, running distance showed a decline in older mice across both strains (p’s < 0.05) (Figures 3C & 3D).

**Figure 3.**
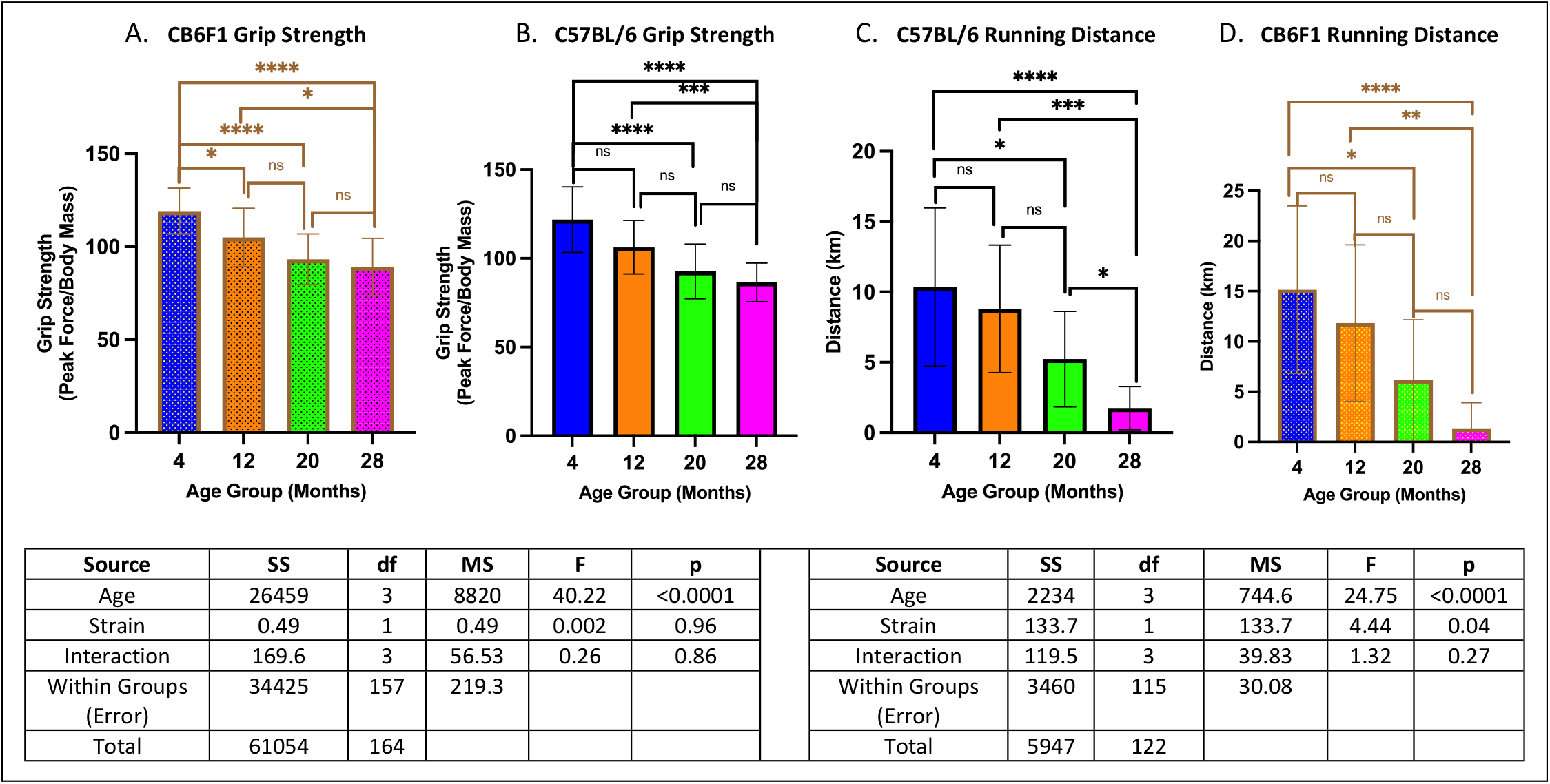
Grip Strength & Running Distance. **A/B**. Both B6 and CB6F1 mice exhibited decreased grip strength with age. **C/D**. Both B6 and CB6F1 mice exhibited decreased running distance with age (*p < 0.05, **p < 0.01, ***p < 0.001, ****p < 0.0001, ns = not significant (p > 0.05), N = 19-27/cohort for grip strength, N = 12-18/cohort for running distance).

### Older B6 mice had higher anxiety compared to CB6F1 Mice

Older B6 mice at the 20- and 28-month cohorts had higher anxiety compared to their CB6F1 counterparts (p’s < 0.05) while no differences were observed between the younger cohorts at 4- and 12-months (p’s > 0.05) (Figure 4).

**Figure 4.**
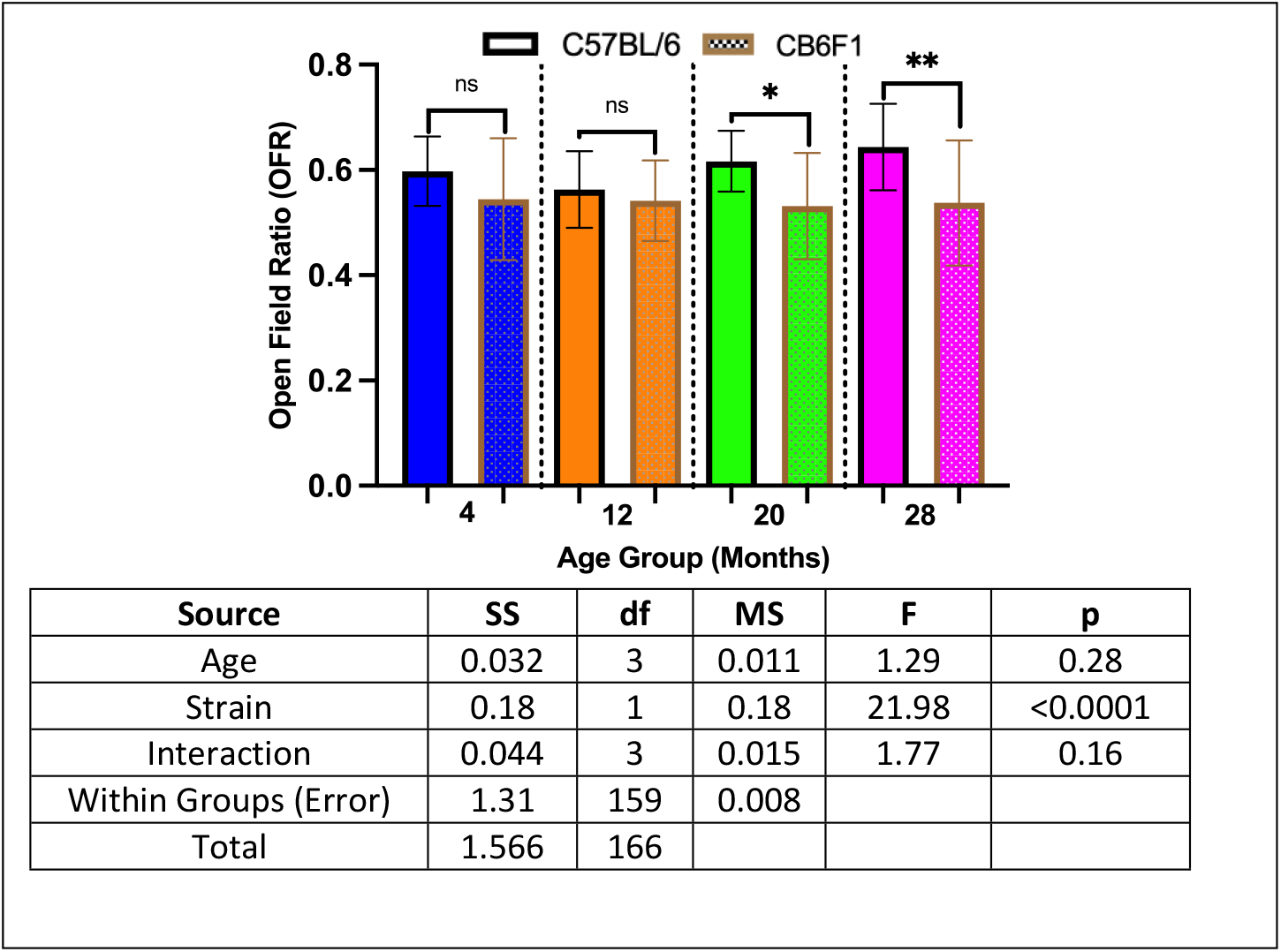
Anxiety Scores. B6 and CB6F1 mice had similar anxiety scores at the 4- and 12-month cohorts although B6 mice had significantly higher anxiety at 20- and 28-months of age compared to CB6F1 mice (*p < 0.05, **p < 0.01, ns = not significant (p > 0.05), N = 19-28/cohort).

### Cataract severity increased in an age-dependent manner in both B6 and CB6F1 mice

Both strains exhibited similar performance across age groups (p’s > 0.05), with cataract severity progressively increasing with age (p’s < 0.01) (Figures 5A & 5B).

**Figure 5.**
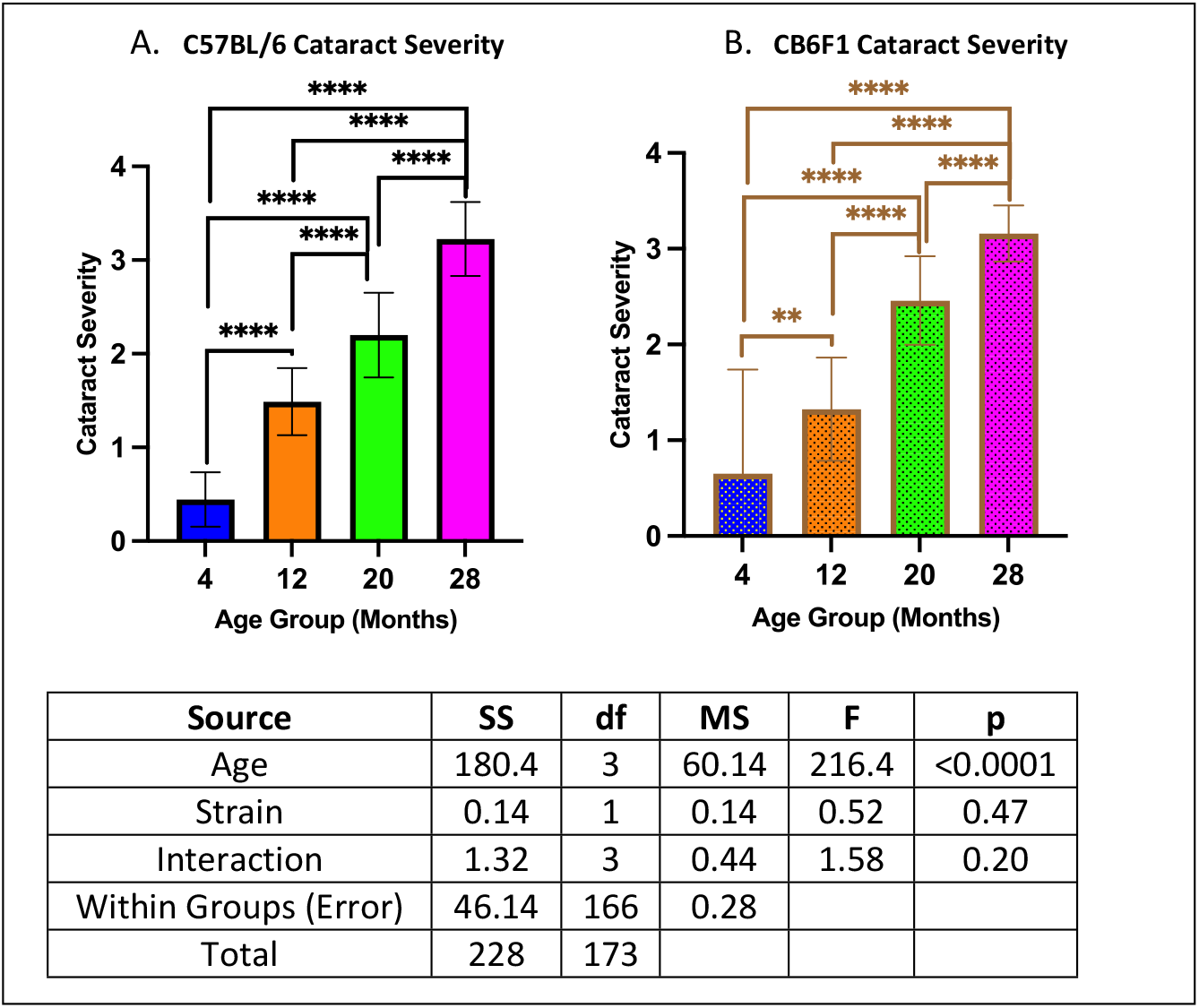
Cataract Severity. **A/B**. Both B6 and CB6F1 mice exhibited greater cataract severity with age (**p < 0.01, ****p < 0.0001, N = 19-28/cohort).

### Metabolic activity was age- and strain-dependent

Significant differences in body fat were observed only between the 4- and 12-month cohorts in both strains (p’s < 0.001) (Figures A & 7B). However, CB6F1 mice consistently exhibited higher body fat levels compared to B6 mice across all age groups (p’s < 0.05) (Figure 7C). Additionally, differences in percent lean body mass were evident in both strains across age groups (p’s < 0.05) (Figure 7D & E), with B6 mice consistently displaying higher percent lean body mass compared to CB6F1 mice across all age groups (p’s < 0.01) (Figure 7F).

**Figure 7.**
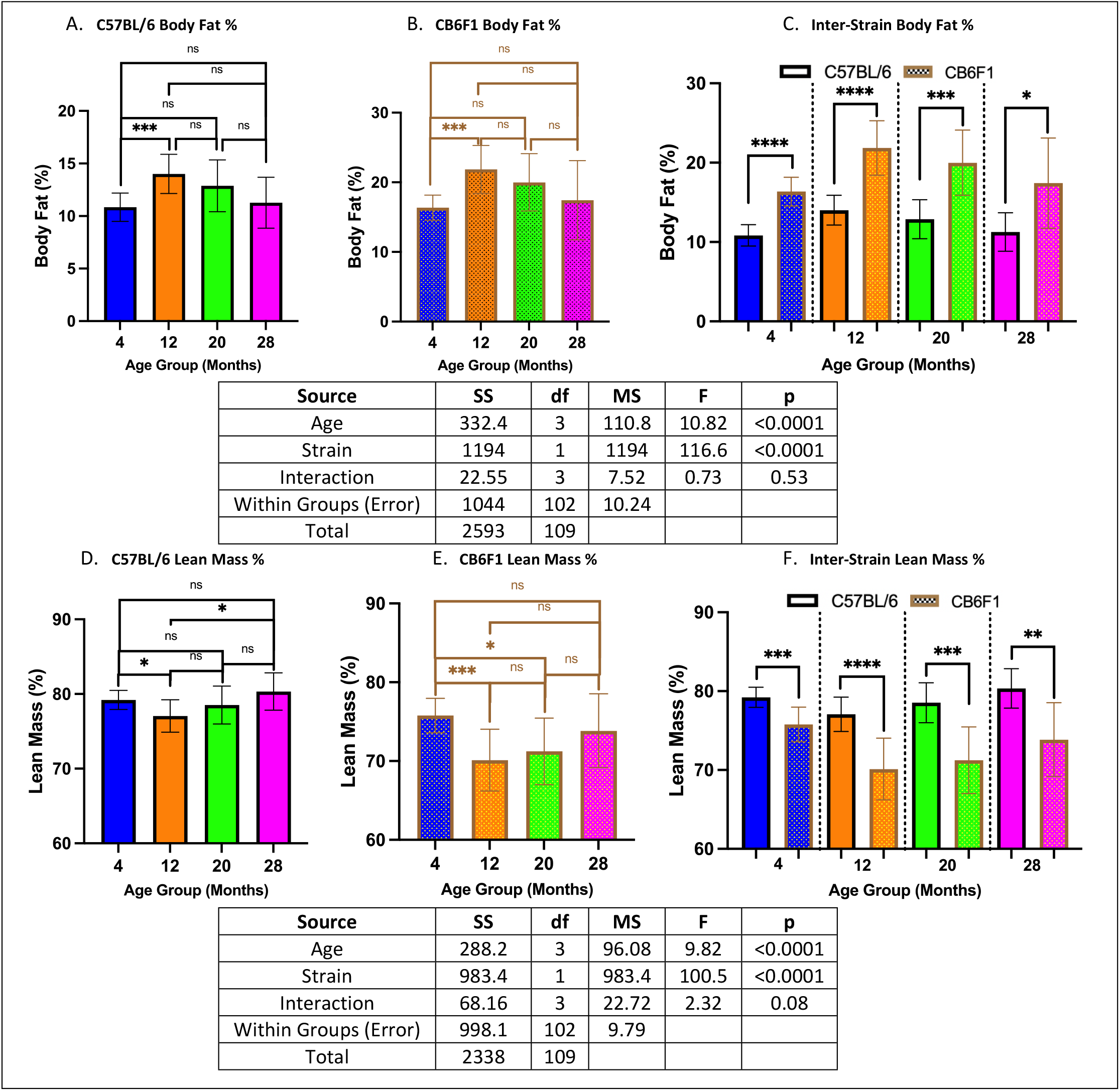
Metabolic Assessment. **A/B**. B6 and CB6F1 mice had relatively stable body fat with only a significant difference between 4- and 12-month cohorts. **C**. Overall, CB6F1 mice had significantly higher body fat than their B6 counterparts across age groups. **D/E**. B6 and CB6F1 mice showed age-related differences in lean body mass. **F**. Overall, B6 mice had significantly higher lean body mass compared to their CB6F1 counterparts across age groups (*p < 0.05, **p < 0.01, ***p < 0.001, ****p < 0.0001, ns = not significant (p > 0.05), N = 10-15/cohort).

### Cardiac functions were age- and strain-dependent

Left ventricular mass index normalized to tibial length (LVMI (tibia)) was used as a measure of left ventricular hypertrophy (LVH). No age-related changes were observed in B6 mice (p’s > 0.05) (Figure 8A), with differences detected only between the 4- and 28-month CB6F1 cohorts (p < 0.05) (Figure 8B). Inter-strain comparisons revealed significantly higher LVMI (tibia) in CB6F1 mice compared to their B6 counterparts (p’s < 0.05) (Figure 8C).

**Figure 8.**
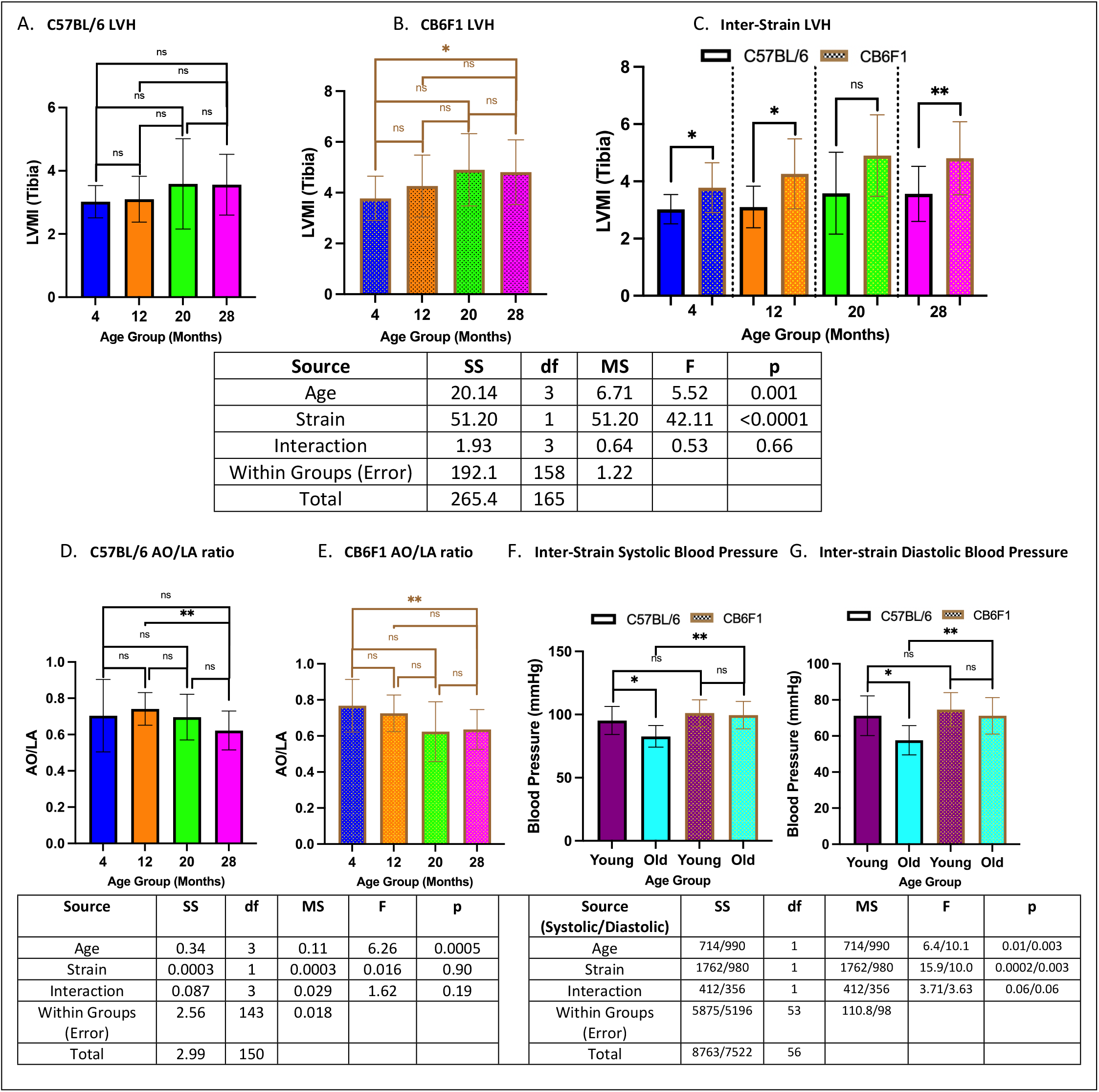
Cardiac function. **A**. B6 mice did not show age-related changes in LVMI (tibia). **B**. 28-month old CB6F1 mice had significantly lower LVMI (tibia) compared to their 4-month old counterparts. **C**. CB6F1 mice had generally higher LVMI (tibia) than B6 mice. **D/E**. Age-related changes in AO/LA ratio were observed in both B6 and CB6F1 mice, with lower ratios in the older age groups. **F/G**. Only B6 mice had age-related decreases in systolic/diastolic blood pressure, while older CB6F1 mice exhibited higher readings compared to their B6 counterparts (*p < 0.05, **p < 0.01, ns = not significant (p > 0.05), N = 18-21/cohort for LVMI (tibia), N = 17-24/cohort for AO/LA ratio, N = 10-16/cohort for blood pressure).

The aortic root to left atrial ratio (AO/LA) served as another measurement of structural heart abnormalities alongside LVMI (tibia). Differences between older and younger cohorts were observed in both B6 and CB6F1 strains, with lower AO/LA ratio detected in the older cohorts (p’s < 0.01) (Figures 8D & 8E).

Additionally, average diastolic and systolic blood pressures were measured as variables of cardiac function. Age-related decreases in both systolic and diastolic blood pressures were observed for B6 mice only (p’s < 0.05) (Figure 8F). However, 20- and 28-month old CB6F1 mice exhibited higher systolic and diastolic blood pressure compared than their B6 counterparts (p’s < 0.05)(Figure 8G). No associations with age or strain were found for measures of left ventricular diastolic dysfunction (i.e., Ea/Aa ratio, IVCT T1, IVRT T3), cardiac reserve (i.e., MPI), or changes in ejection fraction (i.e., FS %).

### Geropathology of the heart, liver, and kidney were age-dependent in both CB6F1 and B6 mice while CB6F1 had more severe lung lesions

There were age-related increases in heart lesion severity in both CB6F1 and B6 strains (p’s < 0.01) (Figures 9A & 9B). Similarly, age-related increases in liver lesion severity were observed in both strains (p’s < 0.05) (Figures 9C & 9D).

**Figure 9.**
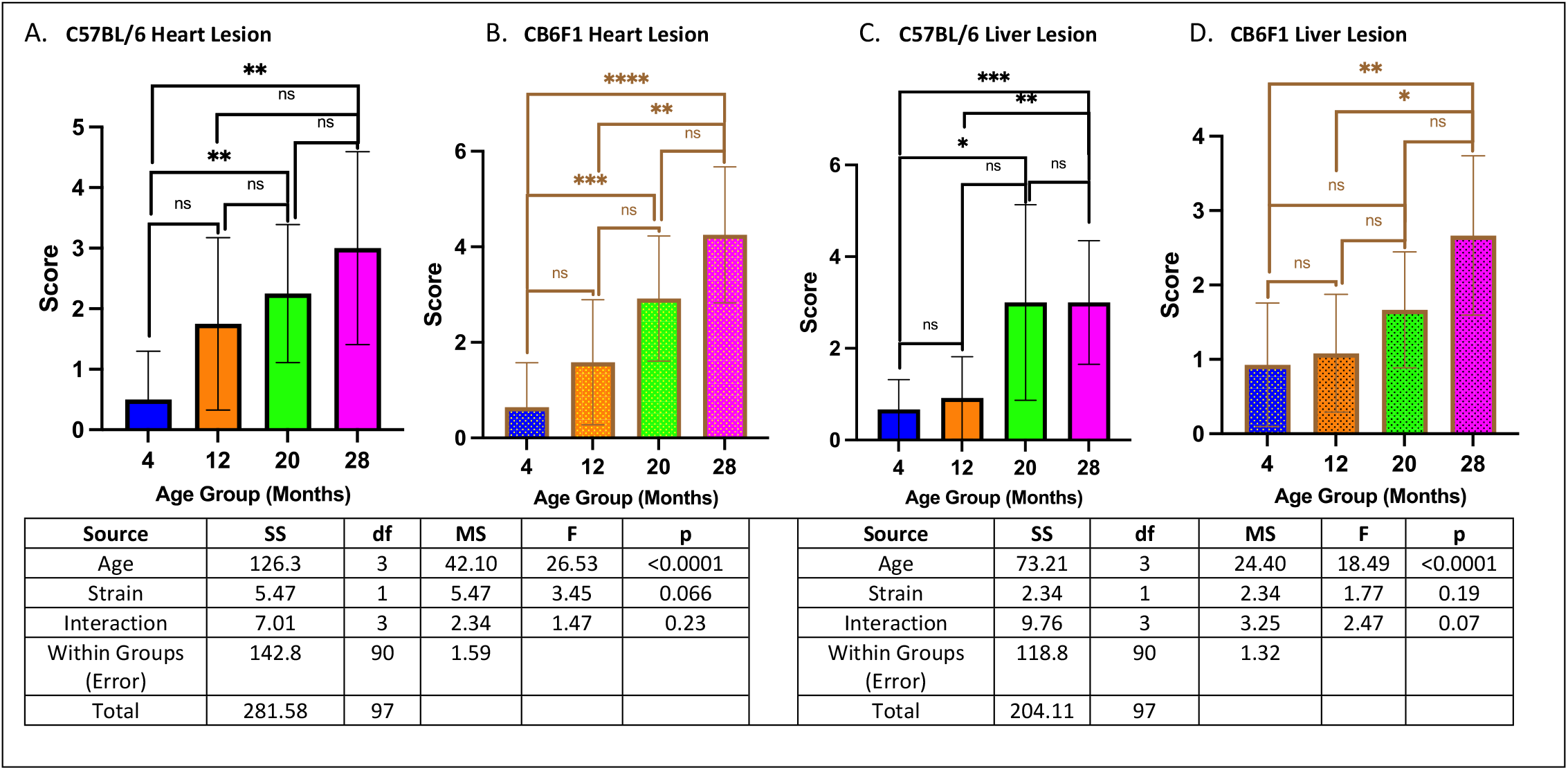
Heart/liver lesion scores. **A/B**. Both B6 and CB6F1 exhibited age-related worsening heart lesion severity. **C/D**. Both B6 and CB6F1 mice exhibited age related worsening liver lesion severity (*p < 0.05, **p < 0.01, ***p < 0.001, ****p < 0.0001, ns = not significant (p > 0.05), N = 12-14/cohort).

Both B6 and CB6F1 mice exhibited increased kidney lesion severity with age (p’s < 0.05) (Figures 10A & 10B), with 20-month-old B6 mice showing more severe kidney lesions than their CB6F1 counterparts (p < 0.05) (Figure 10C). Additionally, there was an age-related increase in lung lesion severity in CB6F1 mice (p’s < 0.05) but not in B6 mice (p’s > 0.05)(Figure 10D & 10E), without any differences between strains (p’s > 0.05)(Figure 10F).

**Figure 10.**
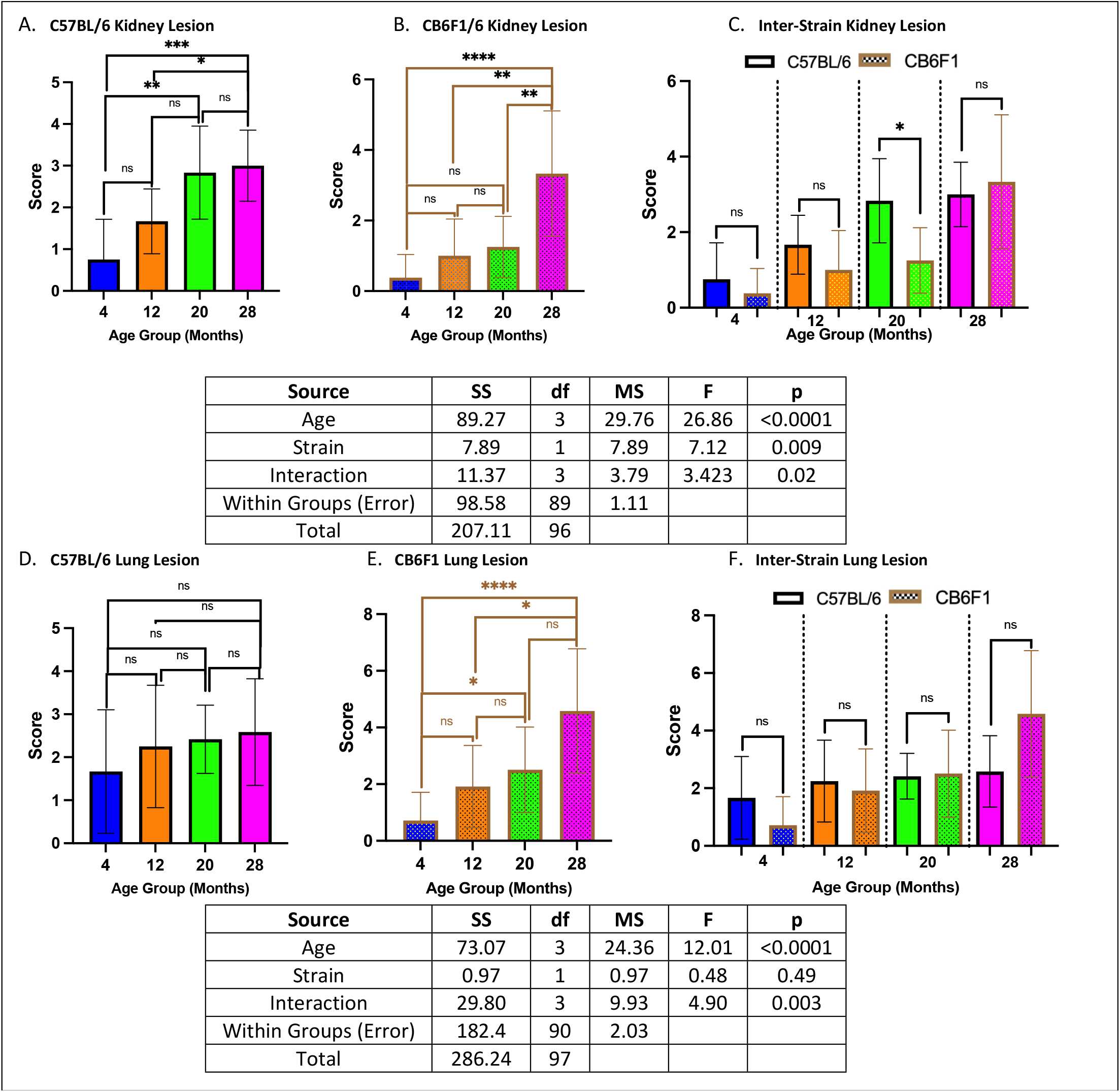
Kidney/lung lesion scores. **A/B**. Both B6 and CB6F1 mice exhibited age-related increases in kidney lesion severity. **C**. Only 20-month-old B6 mice had significantly higher degrees of kidney lesion compared to their age-adjusted CB6F1 counterparts. **D**. B6 mice did not have significant age-related differences in lung lesion severity. **E**. CB6F1 mice had age-related increases in lung lesion severity. **F**. B6 and CB6F1 mice had similar amounts of lung lesion severity across all age groups (*p < 0.05, **p < 0.01, ***p < 0.001, ****p < 0.0001, ns = not significant (p > 0.05), N = 12-14/cohort).

Overall comparison of age- and strain-related differences from the results are summarized in Table 1.

**Table 1.** Summary of results. “+”: evidence of age-related changes, “-”: no evidence of age-related changes, “>“: better performance in a particular strain, “<“: worse performance in a particular strain, “=“: no differences in severity of performance between strains, “↑”: increased severity compared to the other strain, “↓”: decreased severity compared to the other strain.

| Table 1. |  | CB6F1 |  | C57BL/6 |  |
| --- | --- | --- | --- | --- | --- |
|  |  | Age | Strain | Age | Strain |
| Behavioral Assays | Rotarod Performance | + | < | + | > |
|  | Radial Water Tread Maze |  |  |  |  |
|  | STM | + | < | - | > |
|  | LTM | + | = | - | = |
|  | Grip Strength | + | = | + | = |
|  | Voluntary Running Distance | + | = | + | = |
|  | Open Field Test |  |  |  |  |
|  | Anxiety | - | < | - | > |
| Phenotypic Assessment | Cataract Severity | + | = | + | = |
|  | Metabolic Activity |  |  |  |  |
|  | Body Fat % | + | ↑ | + | ↓ |
|  | Lean Body Mass % | + | ↓ | + | ↑ |
|  | Cardiac Functions |  |  |  |  |
|  | LVMi (tibia) | + | ↑ | - | ↓ |
|  | AO/LA ratio | + | = | + | = |
|  | Average Diastolic BP | - | ↑ | + | ↓ |
|  | Average Systolic BP | - | ↑ | + | ↓ |
|  | Ea/Aa ratio | - | = | - | = |
|  | IVCT T1 | - | = | - | = |
|  | IVRT T3 | - | = | - | = |
|  | MPI | - | = | - | = |
|  | FS% | - | = | - | = |
| Geropathological Lesion Severity | Heart | + | = | + | = |
|  | Liver | + | = | + | = |
|  | Kidney | + | ↓ | + | ↑ |
|  | Lung | + | = | - | = |

## Discussion

Analysis of inter-strain differences and similarities of behavioral, physiological, and geropathological assays between CB6F1 and C57BL/6 (B6) mice revealed insights into the relationships between aging and strain-specificity.

As a characteristic of aging across species is the decline in muscle function, evident in reduced strength and muscle mass (20-22), the ability to maintain neuro-muscular performance is pivotal for extending health-span during aging. To obtain a comprehensive profile of muscular ability while minimizing confounding variables such as motivation and sensory deficits, we assessed rotarod performance, grip strength, and wheel running tasks. Both 12- and 20-month-old B6 mice outperformed CB6F1 counterparts on the rotarod task despite age-related performance decline across tasks. While cataract formation, a well-documented consequence of aging (23) shown to increase in severity with age in both strains, may be partially attributable due to the obstruction in vision, because no significant inter-strain differences were observed, differences in rotarod performance are likely to be caused by another variable. CB6F1 mice exhibited higher body fat and lower lean mass percentages, potentially impacting their ability to maintain balance on the rotating rod. Although metabolic differences have been documented (9, 24-26), variations in maternal care could also contribute to observed disparities. Corder et al., 2023 utilized natural cross-fostering (i.e., trio breeding scheme of one sire and two dams per cage) rather than the typical littermate controls, to mitigate environmental and parental care discrepancies. While differences in metabolic activity were nullified, CB6F1 mice still showed inferior rotarod performance compared to B6 (6), signifying that body composition does not confound rotarod performance. Considering the role of brain structures in motor control may be a promising alternative. Gandhay et al., 2023 identified age-related flattening of the hippocampus and pons in CB6F1 mice (4). Differences in age-related atrophy of the pons, which are involved in sensory processing and motor control, could be pivotal in explaining the inferior motor skills of older CB6F1 mice compared to B6. Further investigation into pons atrophy may elucidate the underlying reasons for these differences.

Given the substantial body of evidence linking anxiety to future cognitive decline (27-30), we conducted an analysis of anxiety-related behaviors using the open-field photobeam testing system to explore potential age-related strain differences. The open-field test (OFT) is a widely accepted method for assessing anxiety in mice, leveraging thigmotaxis, an instinctual tendency to avoid open spaces and intensive light due to predation vulnerability (31). This behavior, deeply rooted in evolutionary survival mechanisms, is associated with challenges in emotional and spatial learning (32), with time spent in peripheral regions serving as a key anxiety parameter, quantified by the open field ratio (OFR). Our study revealed that 20- and 28-month-old B6 mice displayed significantly higher OFR compared to CB6F1 mice, consistent with a prior study from the same dataset showing lower overall movement in CB6F1 mice (33), indicating heightened anxiety in older B6 mice relative to their CB6F1 counterparts. While our findings align with previous observations of an inverse relationship between anxiety-like behavior and activity levels (34-35), they diverge from existing literature suggesting lower baseline levels of anxiety in B6 mice without clear rationale (36-37). Additionally, CB6F1 mice exhibited age-related declines in both short-term and long-term memory (STM & LTM) while B6 mice did not, despite evidence suggesting independence between anxiety and memory impairment (33). Given documented instances of memory impairment coinciding with anxiety symptoms (38-41), an unexplored variable in our study worth investigating further is the rate of adult neurogenesis (AN) between the two strains, which is shown to independently affect anxiety-like behaviors and cognition in rodents (42-44). AN, the ongoing generation of neurons in specific brain regions like the ventral dentate gyrus (DG) of the hippocampus beyond early brain development stages (45), has been linked to anxiety-related behaviors and cognitive functions (31,46-48). Therefore, future studies should delve into these variables to unravel the intricate relationship between anxiety and cognition, particularly regarding potential strain disparities.

Both B6 and CB6F1 mice exhibited age-related cardiac dysfunction. Left ventricular hypertrophy (LVH) showed significant differences across all age groups, becoming more severe in 28-month-old mice, particularly in the CB6F1 strain, which exhibited a higher incidence and more severe cardiac dysfunction. This observation is supported by histopathological findings, revealing age-related cardiac lesions such as thickening and fibrosis of the tricuspid and pulmonary valves in CB6F1, which were absent in B6 mice (unpublished findings). The aortic root to left atrial (AO/LA) ratio remained similar in both strains with aging, indicating age-related vessel thickening. While average diastolic and systolic blood pressures increased with age in both strains, older CB6F1 mice showed a greater blood pressure elevation than B6 mice. No significant associations were found between age or strain and other cardiac changes and functions (i.e., Ea/Aa ratio, IVCT T1, IVRT T3, MPI, FS%).

The cardiac aging process observed in mice closely mirrors human cardiac aging, characterized by cardiac hypertrophy, fibrosis, diastolic dysfunction, increased blood pressure, reduced functional reserve, and adaptive capacity to stress (49). This parallel is crucial for understanding human cardiac aging, as the observed reduction in function contributes to the development of heart failure. However, selecting the appropriate animal model, such as rodents, and determining the relevant strain for specific studies and desired lesions is essential to avoid misinterpretation. Similar to humans, genetic background and family history play a significant role in cardiac function and heart health history, emphasizing the importance of careful consideration when using animal models for translational research. In this regard, CB6F1 mice would appear to be an excellent model to study age-related conditions involving function of the left ventricle.

Geropathology assessment of age-related lesions can provide useful information as to how drug treatment can effectively target organs and specific lesions within organs (3). Data from this study showed an age-related increase in lesion scores of the heart, liver, and kidney in both CB6F1 and B6 mice, as expected. The heart lesion scores reflected a similar increase in left ventricular dysfunction as already discussed. We did not conduct any specific liver function tests and found no associations between liver scores and metabolic data in either strain (p > 0.05). Since the liver is involved in a number of metabolic activities, additional consideration of the relationship between liver lesions and metabolic activities is warranted.

There was an age-related increase in lesion scores of the kidney that was more severe in B6 mice compared to CB6F1. This is visually seen with increasing age in a graphic overview (Figure 11), which provides an example of how composite lesion scores (CLS) can distinguish lesion severity differences in the same organ from mice of the same age but of differing genetic backgrounds. Clearly, it can be seen that B6 mice attain a specific composite lesion score much earlier than CB6F1 mice. For example, at 24 months of age CB6F1 mice have a CLS similar to B6 mice at 16 months of age suggesting that the CLS system can be used to indicate the degree of pathological aging, a so-called “pathobiological clock”.

**Figure 11.**
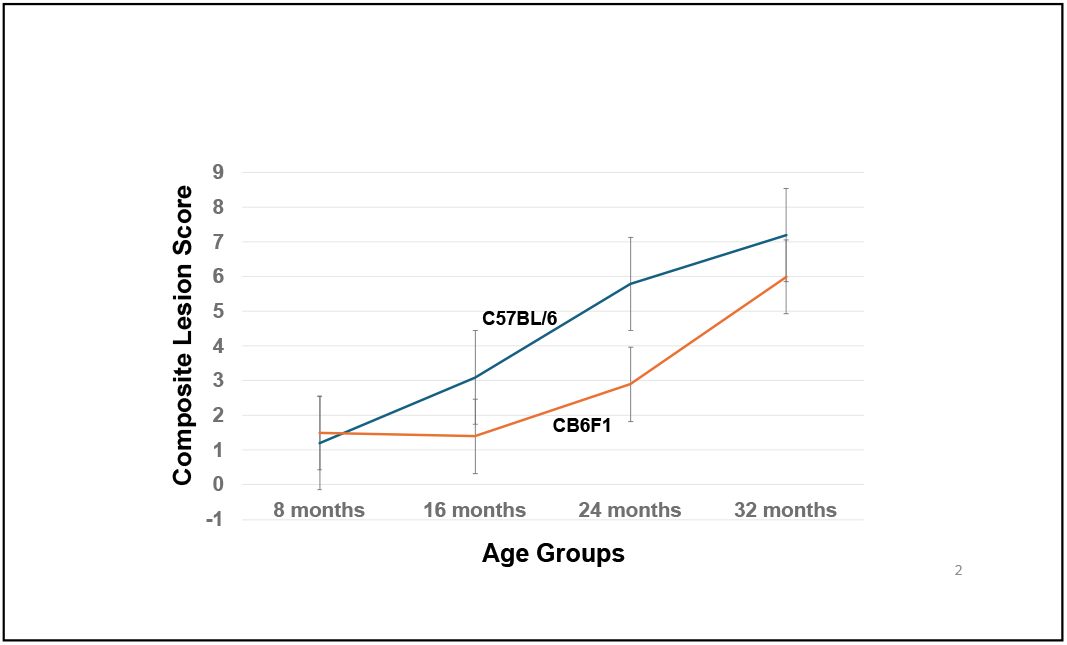
The kidney ages more rapidly in C57BL/6 mice compared to CB6F1 mice. CB6F1 mice at 24 months have kidneys with a pathobiological age of 16 months.

The age-related increase in lesion scores of the lungs in CB6F1 mice, but not B6 mice, is another example of how CLSs can distinguish differences in organ aging. We have previously reported that CB6F1 mice have a rather high incidence of primary lung tumors (50), which could be associated with the progression of lesion burden with increasing age in this strain, whereas B6 mice have very few primary lung tumors.

In summary, the genetic background of the two mouse strains influenced results of bioassays in an age-dependent manner. Therefore, it is imperative to recognize that different strains of mice may yield diverse data in preclinical studies and would need to be interpreted individually for translational applications.

## Acknowledgement

This study was supported by National Institute on Aging grant R01 AG057381.

